# Primordial germ cells adjust their protrusion type while migrating in different tissue contexts *in vivo*

**DOI:** 10.1101/2022.11.02.514858

**Authors:** Lukasz Truszkowski, Dilek Batur, Hongyan Long, Katsiaryna Tarbashevich, Bart E. Vos, Britta Trappmann, Erez Raz

## Abstract

In both physiological processes and disease contexts, migrating cells have the ability to adapt to conditions in their environment. As an *in vivo* model for this process, we use zebrafish primordial germ cells that migrate throughout the developing embryo. When migrating within an ectodermal environment, the germ cells form fewer and smaller blebs as compared with their behavior within mesodermal environment. We find that cortical tension of neighboring cells is a parameter that affects blebbing frequency. Interestingly, the change in blebbing activity is accompanied by the formation of more actin-rich protrusions. These alterations in cell behavior that correlate with changes in RhoA activity could allow the cells to maintain dynamic motility parameters, such as migration speed and track straightness, in different settings. In addition, we find that the polarity of the cells can be affected by stiff structures positioned in their migration path.

## Introduction

Primordial germ cells (PGCs) are typically specified away from the region where the gonad develops, such that they must migrate toward a region in the embryo where they associate with cells of mesodermal origin to form the gonad (Barton et al., 2016; Grimaldi and Raz, 2020; Richardson and Lehmann, 2010).

During their migration, PGCs encounter multiple types of cells and tissues. This is especially evident in zebrafish, where PGCs are specified at four locations that are randomly oriented relative to the embryonic axis (Weidinger et al., 1999). Starting from the positions where they originate, PGCs migrate through the embryo as the tissues around them undergo processes of differentiation and morphogenesis. Accordingly, to reach their target, PGCs have to move within tissues that exhibit different biophysical properties.

The migration of zebrafish PGCs is guided by the chemokine Cxcl12a, which binds the receptor Cxcr4b (Doitsidou et al., 2002). Interestingly, in the absence of the guidance cue or its receptor, the PGCs remain motile and migrate non-directionally within the whole embryo (Doitsidou et al., 2002; Gross-Thebing et al., 2020).

Zebrafish PGCs perform ameboid motility, which is characterized by extensive cell body deformations, low substrate attachment, and generation of actin-based pseudopods or hydrostatic pressure-powered blebs (Paluch and Raz, 2013; Yamada and Sixt, 2019; Charras et al., 2006; Schick and Raz, 2022). A range of studies conducted mainly in *in vitro* settings show that cells can alter their protrusion type or migration mode when located in different 2D and 3D environments (Lämmermann and Germain, 2014; Liu et al., 2015; Talkenberger et al., 2017; te Boekhorst et al., 2016). Here we use zebrafish PGCs as an accessible *in vivo* model for exploring how amoeboid cells respond to changes in the properties of their environment.

We show that PGCs migrate efficiently in different cellular contexts and respond to differences in biophysical properties of their environment by adjusting the protrusion type they produce. We also highlight the cortex tension of surrounding cells as a parameter influencing the protrusive activity.

## Results and Discussion

### Migration of PGCs within different embryonic germ layers

PGCs lacking the guidance receptor Cxcr4b can be found within ectodermal, mesodermal and endodermal tissues (Fig. S1A, Gross-Thebing et al., 2020). These different germ layers present PGCs with environments that have distinct biophysical properties (Krieg et al., 2008), allowing us to study the behavior of the cells in different tissue contexts within the live animal. To ensure uniform conditions around the Cxcr4b-depleted PGCs, we generated single-germ-layer embryos according to previously established protocols (Krieg et al., 2008) (Fig. 1A-D, Fig. S1B). Importantly, these experimental conditions do not affect the identity of PGCs, as determined by the expression levels of germ cell-specific RNA markers (Fig. S1C-D), in line with studies demonstrating the robustness of the PGC fate in face of somatic differentiation cues (Gross-Thebing et al., 2017; Strome and Updike, 2015).

**Figure 1.**
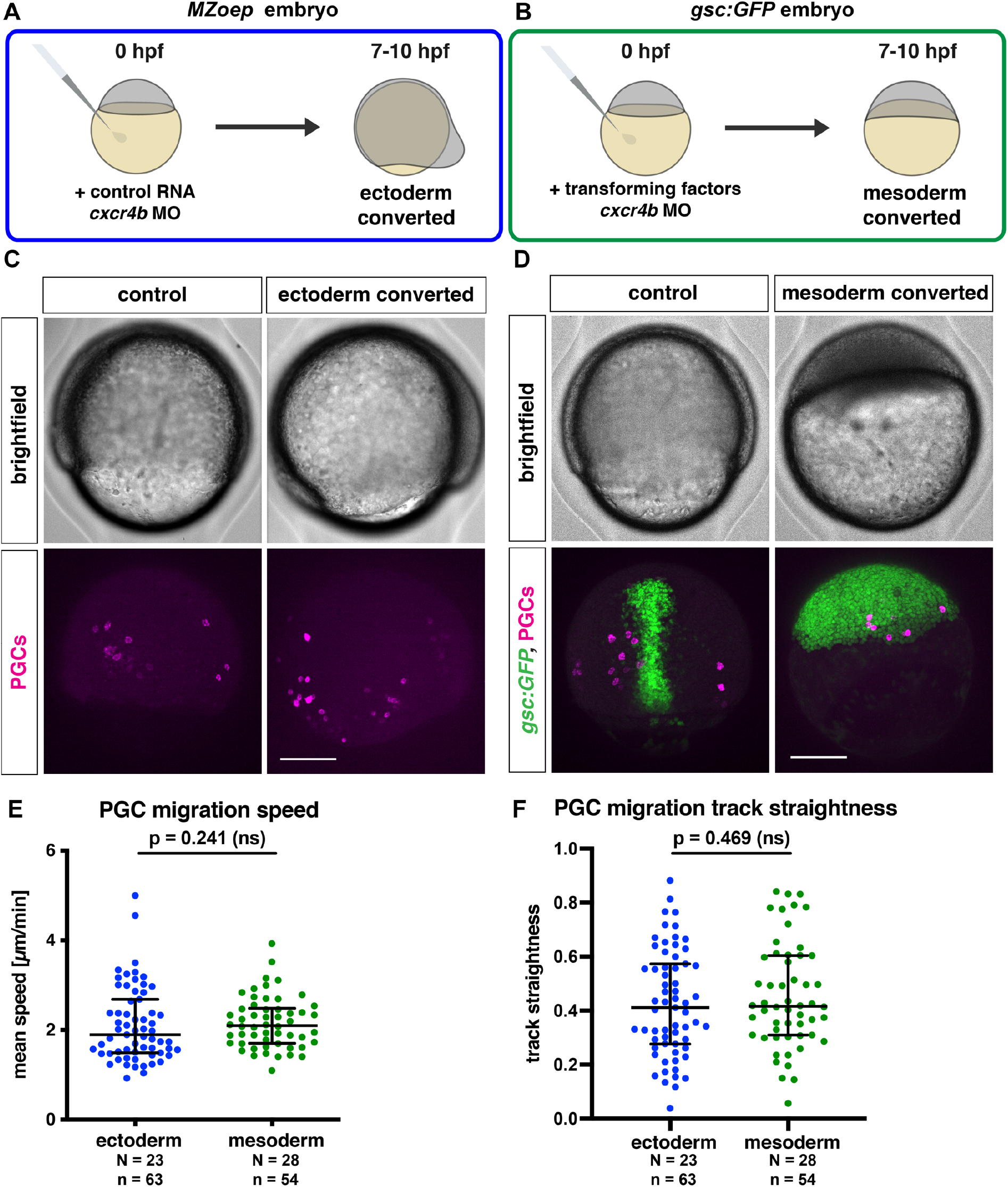
Migration of PGCs within ectodermal and mesodermal tissues. (A), Schematic depiction of the generation of embryos composed of ectodermal cells. In maternal-zygotic *one eye pinhead* mutant embryos (*MZoep*), all of the somatic cells develop into ectoderm, with Cxcr4b expression inhibited by injection of *cxcr4b* morpholino. (B), Conversion of embryonic cells into mesoderm is achieved by co-injection of RNA encoding for a Nodal ligand (*cyclops*) and morpholino directed against the RNA encoding for the transcription factor Sox32 (together termed “transforming factors”). (C), Lateral view of control and ectoderm-converted embryos (upper panels), with the PGCs labeled (magenta, lower panels). (D), Dorsal view showing control and mesoderm-converted embryos (upper panels) with the PGCs labeled (magenta) and the expression of GFP driven by the *goosecoid* promoter (green) in the corresponding embryos (lower panels). Scale bars in C and D, 200 μm. (E-F), Migration speed (E) and track straightness (F) of PGCs migrating within ectodermal and mesodermal tissues. Lines show median value +/− interquartile range (IQR). For speed comparison (E), Mann-Whitney test was performed; for straightness comparison (F) two-tailed t-test was performed. N and n represent numbers of embryos and cells, respectively.

As a first step in characterizing the dynamic behavior of PGCs located within different germ layers, we monitored their migration speed and track straightness. Interestingly, we found these parameters to be similar for cells located within the ectoderm and the mesoderm, despite the previously described differences in tissue properties (Fig. 1E-F).

### The effect of the cellular environment on bleb dynamics

Zebrafish PGCs primarily generate bleb-type rapidly inflating round actin-free protrusions that facilitate the movement of the cell (Blaser et al., 2006; Goudarzi et al., 2017; Olguin-Olguin et al., 2021). To determine whether features of the cellular environment influence the protrusive activity, we monitored the frequency of bleb formation and the size of blebs. Interestingly, PGCs migrating within the ectodermal tissue formed fewer and smaller blebs, as compared with those formed by cells located within the mesodermal tissue (Fig. 2A-C). These results were further supported by analyzing blebbing in wild-type PGCs transplanted into either ectoderm-converted or control embryos. Here as well, fewer and smaller blebs were generated by PGCs transplanted into the ectodermal environment as compared with controls (Fig. S2).

**Figure 2.**
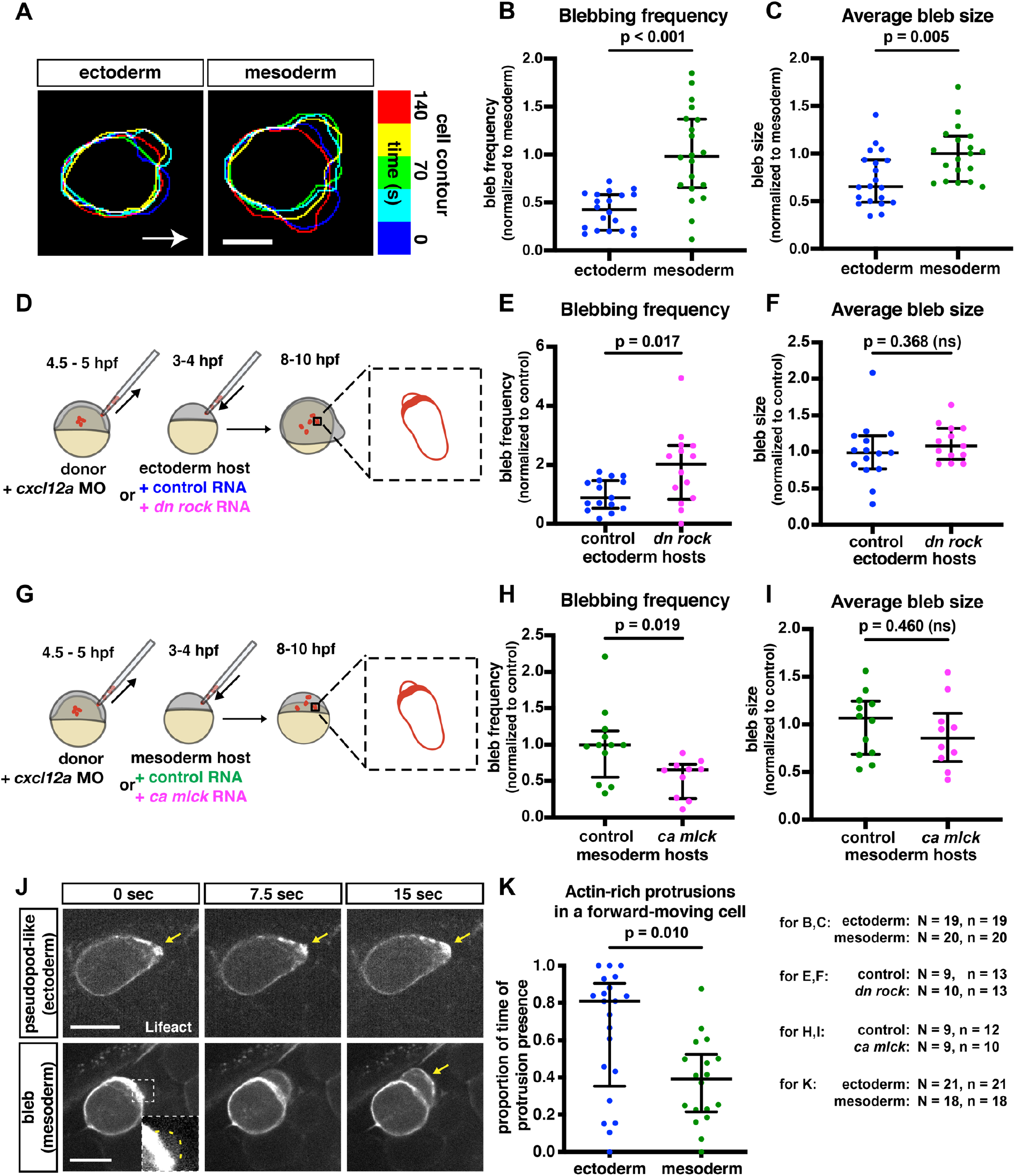
Alterations in protrusive behavior in PGCs located within different environments. (A), Overlay of cell contours over time of representative PGCs migrating in ectodermal and mesodermal tissues. Contours were aligned to the back of the cell to correct for forward movement. Arrow indicates the direction of migration. Scale bar, 10 μm. (B-C), Blebbing frequency (B) and average bleb size (C) of PGCs moving within converted embryos. For bleb frequency comparison (B), Mann-Whitney test was performed; for average bleb size (C) two-tailed t-test was performed. (D), The PGC transplantation experiment scheme. Labeled wild-type PGCs were transplanted into ectoderm-converted embryos that were injected at the 1-cell stage with either control RNA or a dominant-negative form of ROCK. 4 to 6 h later, blebs formed by the transplanted PGCs were analyzed. (G), A similar transplantation was performed into mesoderm-converted embryos that expressed either control RNA or RNA encoding for a constitutively active form of MLCK. (E-F,H-I), Bleb formation frequency (E, H) and average bleb size (F, I) produced by PGCs transplanted into the different environments. Two-tailed t-tests were performed. (J), Snapshots of migrating PGCs. Arrows point at actin enrichment at the edge of protrusion. Dashed box presents a zoomed-in, contrast-adjusted view on the early-stage bleb, with the yellow dashed line outlining the bleb contour. Scale bars, 15 μm. (K), Proportion of time actin-rich protrusions are present at the front of a forward-moving cell. Mann-Whitney test was used. For all the plots, lines show median +/− IQR. N and n at the bottom right represent numbers of embryos and cells, respectively.

Since blebs are formed more frequently in cells migrating under confinement (Liu et al., 2015; Ruprecht et al., 2015), we examined the density of the nuclei within converted embryos and found that it was not different between ectodermal and mesodermal tissues (Fig. S3).

Relative to the other germ layers, ectodermal cells possess higher cortex tension, which is primarily generated by myosin contractility (Krieg et al., 2008). To examine whether this feature affects the behavior of the PGCs, we reduced contractility in ectodermal cells (Fig. 2D) and transplanted PGCs into the manipulated environment. Interestingly, transplanted PGCs that migrated within the lower-contractility ectodermal tissue formed blebs more frequently (Fig. 2E), without changing the bleb size (Fig. 2F). Conversely, PGCs transplanted into mesodermal tissue in which contractility was increased (Fig. 2G) formed fewer blebs (Fig. 2H) with no change in bleb size (Fig. 2I). These observations suggest that the cortex tension in cells surrounding the PGCs is an important parameter that determines how often blebs form. However, while the cortex tension of cells in the environment influences the frequency of bleb initiation, this parameter does not affect bleb expansion. This suggests that in addition to cortical tension, other parameters influence blebbing in these settings. Consistent with the results presented above, the migration speed and track straightness were similar for cells migrating within control and ectodermal environment in which contractility was lowered (Fig. S4).

### Alterations in protrusion types

Since PGCs mainly extend blebs (Olguin-Olguin et al., 2021), it is surprising that ectoderm-residing cells that generate fewer and smaller blebs migrate similarly to mesoderm-residing PGCs concerning speed and track straightness (Fig 1E-F, Fig. S4). To gain further insight into the cellular mechanisms that govern protrusion formation in both cases, we examined the distribution of actin within the protrusions. Interestingly, during active migration phases, PGCs located within ectodermal tissue formed actin-rich protrusions more often than cells located within mesodermal tissue (Fig. 2J-K). In those cases, actin was continuously present at the constantly advancing leading edge (Movie 1, Fig. 2J upper panels), in contrast with the temporary lack of actin at the front of the intermittently forming, rapidly expanding bleb (Movie 2, Fig. 2J lower panels). Interestingly, we found that protrusions characterized by elevated levels of actin at their leading edge form a three-dimensional structure, as blebs do (Movie 3). Based on these characteristics, we conclude that when located within the ectodermal environment, PGCs adjust their protrusion type and form more actin-driven pseudopod-type protrusions. Importantly, as measured in fibroblasts, an actin cortex-bound membrane exhibits 10 times higher tension than blebs that lack actin underneath their membrane (Tinevez et al., 2009). The increased tension in the actin-rich protrusion could make migration more effective within the ectoderm. In the context of the wild-type embryo, this shift could help PGCs exit the ectoderm and reach the developing gonad, which is a mesodermal derivative.

### The molecular organization of migrating PGCs in different cellular environments

Next, we examined parameters that could account for the observed alterations in the protrusion types. To this end, we determined the activation state of RhoA and Rac1, Rho-GTPases that control contractility and actin polymerization (Ridley, 2015), and are important for PGC migration (Kardash et al., 2010; Olguin-Olguin et al., 2021). Employing FRET-based sensors, we found that germ cells located within ectodermal tissue exhibited a mild increase in RhoA activity, which was similar at the front and the back of the cell, while Rac1 activity was not altered (Fig. 3A-B, Fig S5). To examine whether the increase in RhoA activity is associated with an elevation in myosin activity, we employed a myosin light chain (MLC) FRET-based sensor (Yamada et al., 2005) and observed no difference in MLC-FRET between PGCs located in ectoderm or mesoderm (Fig. 3C). Similarly, actin retrograde flow, which is controlled by actin polymerization, adhesion and actomyosin contraction (Renkawitz et al., 2009), was not different between the two conditions (Fig. 3D-E).

**Figure 3.**
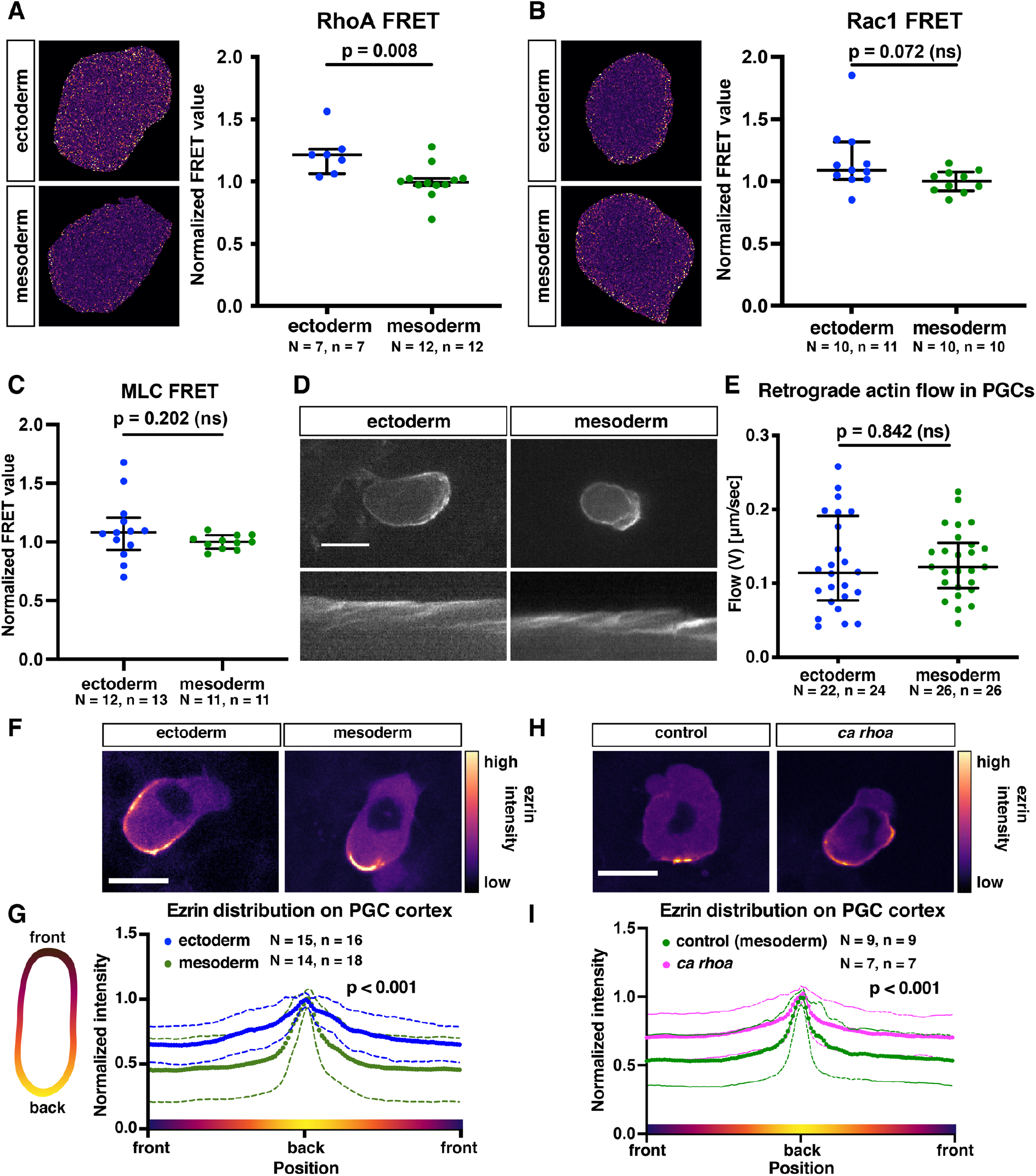
The role of Rho GTPases in controlling protrusion type formation. (A-B), Normalized whole cell FRET-based activity measurement of RhoA (A) and Rac1 (B). Lines show median +/− IQR. Two-tailed t-test was performed. Mann-Whitney test was performed. (C) Normalized whole cell FRET-based myosin light chain phosphorylation levels. Lines show median +/− IQR. Two-tailed t-test was performed. (D), Representative images of actin flow in PGCs migrating within ectodermal and mesodermal tissues. Kymographs derived from the middle of the cell. Scale bar, 15 μm. (E), Retrograde actin flow speed (V). Lines show median +/− IQR. Two tailed t-test was performed. (F, H), Representative examples of ezrin-YPet distribution in PGCs residing within converted embryos (F), or in PGCs expressing control RNA or low levels of RNA encoding for constitutively active form of RhoA (H). Scale bars, 15 μm. (G,I), Schematic representation of cortical ezrin distribution around the cell perimeter (left). Cortical ezrin signal was masked and sliced in angular manner, and the results are presented in the graphs. For each cell, ezrin intensity was normalized to the peak intensity in the cell. Colored dots and solid lines represent the means, while dashed lines represent the standard deviations. Kolmogorov-Smirnov test was performed to compare frequency distributions. N and n represent numbers of embryos and cells, respectively.

Relevant for this work, in addition to increasing contractility, RhoA activity can promote ezrin, radixin and moesin (ERM) activation (Bagci et al., 2020; Hebert et al., 2008; Matsui et al., 1999), which could inhibit blebbing by linking the membrane to the underlying cortex and endoplasmic reticulum (Charras et al., 2006; Olguin-Olguin et al., 2021). To investigate the possible involvement of ezrin in the altered cell behavior, we examined its distribution around the cell perimeter and found it to be more evenly distributed in PGCs migrating within the ectoderm (Fig. 3F-G). Importantly, a mild increase of RhoA activity in PGCs migrating within the mesoderm resulted in distribution of ezrin that is similar to that observed in ectoderm-residing PGCs (Fig. 3H-I).

### The effect of protrusion type change on migration

As presented above, PGC migration speed and track straightness were similar within different environments (Fig. 1E-F, Fig. S4). It could thus be that protrusion-type shifts reflect adaptations to environment properties, maintaining migration parameters. To test this hypothesis, we examined in what way increasing the frequency of bleb formation affects PGC migration. To this end, we expressed in PGCs low levels of a dominant negative version of Rac1, which led to increased bleb formation (Vidali et al., 2006, Fig. 4A), presumably by reducing the level of actin at the cell cortex. Interestingly, when migrating within ectodermal environment, such PGCs exhibited a strong reduction in migration speed (47%) and track straightness as compared with control cells (Fig. 4B). This effect was less pronounced in PGCs that were treated in the same way and migrated within mesodermal cells (25%, Fig. 4C). These findings suggest that interfering with the ability of germ cells to adjust their protrusion type in specific environments reduces their migration efficiency.

**Figure 4.**
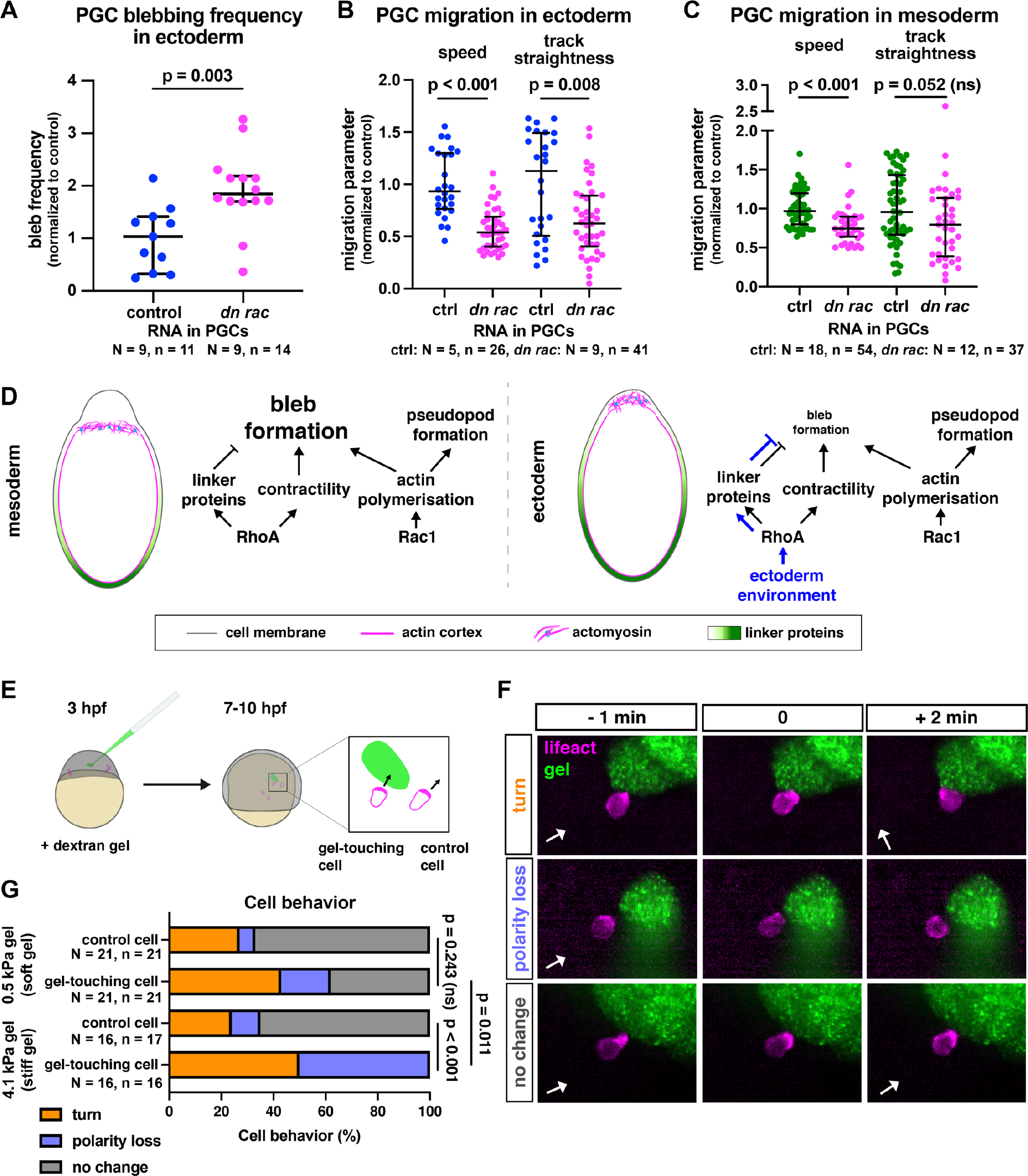
Effect of blebbing and physical barriers on PGC migration and polarity. (A), Blebbing frequency in PGCs that reside within ectodermal tissues and express control RNA or RNA encoding for dominant-negative version of Rac1. Two-tailed t-test was performed. (B-C) Speed and track straightness of PGCs expressing control RNA or a dominant-negative version of Rac1, migrating among converted ectodermal cells (B) or mesodermal cells (C). Mann-Whitney test was performed for track straightness comparison in (B) and both comparisons in (C), two-tailed t-test was performed for speed comparison in (B). For all the plots, lines show median +/− IQR. (D), A model illustrating molecular interactions that lead to the protrusion-type shift in different environments. (E), Illustration of the gel injection experiment. Mutant *cxcr4b^-/-^* embryos were injected with a dextran gel of 0.5 kPa or 4.1 kPa stiffness at 3 hpf and were imaged 4 h later. As cells migrate non-directionally, some stochastically come in contact with the gel (left cell in magnified box). Alterations in actin distribution were then monitored in cells that came into contact with the gel and in neighboring PGCs that did not encounter the gel (right cell in magnified box). The comparison between the PGCs was performed to control for the inherent periodic loss of polarity observed in PGCs (tumbling, Reichman-Fried et al., 2004). (F), Examples of PGCs’ (lifeact in magenta) reactions to the gel (green). Behaviors were scored based on the change in the polarity of the actin-rich front. The white arrows indicate the direction of the back-front axis of the cell before the interaction with the gel. (G), Quantification of cell behaviors. For statistical tests, Fisher’s exact test was performed (comparing the combined turn and loss of polarity behaviors to no change in polarity). Bonferroni correction for multiple testing was applied. N and n represent numbers of embryos and cells, respectively.

### Plasticity in PGC migration – a model

Based on our results, we present a model that accounts for the alterations in protrusion types we observed in different cellular environments (Fig. 4D). In non-manipulated embryos, PGCs migrate within and in close proximity to the mesoderm, where they more readily form blebs (84% blebs, 16% pseudopods, Olguin-Olguin et al., 2021). In response to interaction with cells that exhibit increased cortical tension (e.g. in ectoderm), PGCs mildly increase their RhoA activity. Such an elevation in RhoA activity in response to an increase in tension in the environment has been described before in other contexts (e.g. Acharya et al., 2018; Gupta et al., 2021). It should be noted that very high levels of RhoA activity lead to stable bleb formation (Liu et al., 2015; Olguin-Olguin et al., 2021; Ruprecht et al., 2015). However, in the context of this work, a moderate elevation of RhoA activity could promote activation of ERM proteins, leading to their broader distribution around the cell perimeter (Bagci et al., 2020). We suggest that upon interaction with cells with higher cortex tension, PGCs strengthen the attachment of the cortex to the cell membrane, which inhibits blebbing (Charras et al., 2006; Goudarzi et al., 2017; Niggli and Rossy, 2008). Under these circumstances, the cells form more actin-based protrusions.

Previous research in the lab demonstrated that the migration of PGCs is affected by the lack of E-cadherin in the environment (Grimaldi et al., 2020). Specifically, a strong uniform reduction of E-cadherin-based adhesion resulted in a reduction in migration-track straightness as a result of an increase in retrograde actin flow. Asymmetrical interaction of PGCs with cells devoid of E-cadherin led to asymmetry in actin retrograde flow such that blebs formed away from the manipulated clones, resulting in turning. Cells in ectoderm-converted embryos express lower levels of E-cadherin as compared with that in mesoderm-converted embryos (Krieg et al., 2008), which we could reproduce in our experimental setup (Fig. S6). However, this difference is not as pronounced as that obtained by Grimaldi et al. (Grimaldi et al., 2020), and the retrograde actin flow and migration track straightness are similar in both environments (Fig. 3E-F, Fig. 1E-F). Thus, while E-cadherin function is required for proper migration track straightness (Grimaldi et al., 2020), the differences between the adhesion levels offered by ectodermal and mesodermal cells are not sufficient for affecting this specific parameter.

At this stage, the precise mechanisms by which germ cells sense the cortex tension of cells in their environment are unknown. We have previously characterized interactions of PGCs with the developing gut (Paksa et al., 2016) and the notochord (Gross-Thebing et al., 2020), as well as with cells lacking E-cadherin (Grimaldi et al., 2020). However, the previously described interactions could potentially involve signals that are not purely physical. To examine the cellular response to cues that are purely physical in nature, we introduced dextran-based hydrogels of different stiffnesses into embryos and monitored the interaction of PGCs with the gels. Here, we consider gel stiffness as a parameter related to the cortex tension of cells the PGCs interact with, which could be perceived in a similar way. In these experiments we found that cells interacting with stiffer gels exhibited more pronounced changes in actin polarity as compared with their response towards softer gels (Fig. 4E-G, Fig. S7, Movies 4-6). Notably, while both gels are non-adhesive and thus could promote a loss-of-adhesion based turning (Grimaldi et al., 2020), they induced very different responses in the migrating cells. Our results thus suggest that the behavior of PGCs *in vivo* can be affected by physical interactions that are independent of cell-cell adhesion or other specific molecular signals, such as repulsive cues. Intriguingly, the response of PGCs to the differences in stiffness appears to be associated with actin-rich protrusions. Specifically, 23 out of 27 cases of responses to contact with the gel involved actin-rich cell front, rather than blebs. These results are consistent with the idea that physical interactions of this kind can involve actin-based pushing forces, or proximity of polymerizing actin to tip of the protrusion (see Gaertner et al., 2022).

We have previously shown that migrating PGCs are extremely robust with respect to their ability to invade and migrate within a wide range of tissues in the developing embryo, allowing all the PGCs to arrive at their target (Gross-Thebing et al., 2020; Weidinger et al., 1999). Here, we found that when located in different environments, PGCs change the types of protrusions they produce, while maintaining their migration speed and track straightness. This is consistent with our previous observation of an almost even distribution of unguided PGCs across all germ layers (Gross-Thebing et al., 2020). The ability to migrate within tissues of different properties and the associated plasticity have been also described for leukocytes and cancer cells that change their migration strategies in different *in vitro* settings (e.g., Lämmermann and Germain, 2014; Tozluoğlu et al., 2013). Our findings obtained in an *in vivo* context are thus likely relevant for migration processes of other cell types that move within tissues of different biophysical properties.

## Materials and methods

### Zebrafish strains and maintenance

The following zebrafish (*Danio rerio*) lines were used (allele names in parentheses, according to ZFIN): wild type (AB), *MZoep^tz257^* (*tdgf1^tz257/tz257^* – Gritsman et al., 1999), *Tg(−1.8gsc:GFP)* (ml1Tg – Doitsidou et al. 2002), *Tg(kop:mCherry-FTASE-UTR-nanos3)* (mu6tg – Tarbashevich et al., 2015), *Tg(kop:EGFP-LIFEACT-UTR-nanos3)* (mu4tg – Hartwig et al., 2014), *Tg(kop:EGFP-FTASE-UTR-nanos3)* (er1Tg – Blaser et al., 2005), *Tg(kop:lifeact-mCherry-utr-nanos3,cryaa:dsred)* (mu118Tg – Grimaldi et al, 2020), *cxcr4b^J1049/J1049^* (t26035 – Knaut et al., 2003). The fish maintenance was supervised by the veterinarian office of the city of Muenster, according to the laws of Germany and the state of North Rhine-Westphalia. For details on the fish used in each experiment, see Table S1.

### Zebrafish embryo injection

Capped mRNAs were transcribed *in vitro* using the mMessageMachine kit (Invitrogen, Waltham, MA, USA). A 1 nl drop containing RNAs and/or morpholinos (hereafter (MO), Genetools, Philomath, OR, USA) was injected into early 1-cell stage embryos, unless specified otherwise. The following mRNAs were injected (internal construct designation in parentheses): *lifeact-mCherry-UTR-nanos3* (D554 – Olguin-Olguin et al 2021), *mCherry-FTASE-UTR-nanos3* (A906 – Hartwig et al., 2014), *gfp-FTASE-UTR-globin* (393 – Boldajipour et al., 2011), *mCherry-FTASE-UTR-globin* (A709 – Kardash et al., 2010), *cyclops-UTR-globin* (B836 – Sampath et al., 1998), *h2a-tagBFP-SV40polyA* (D846 – Compagnon et al., 2014), *RacFRET-UTR-nanos3* (A422 – Kardash et al., 2010), *RhoAFRET-UTR-nanos3* (A676 – Kardash et al., 2010), *MLCFRET-UTR-nanos3* (A559 – Blaser et al., 2006), *tdgf1-UTR-globin* (071 – Zhang et al., 1998), *ezrin-YPET-UTR-nanos3* (D025 – Olguin-Olguin et al., 2021), *dn-rock-UTR-globin* (E515 – this work), *ca-mlck-UTR-globin* (E474 – this work), *ca-rhoa-UTR-nanos* (B282 – Kardash et al., 2010), *dn-rac-UTR-nos* (482 – Olguin-Olguin et al., 2021), *mCherry-h2b-UTR-globin* (B325 – Paksa et al., 2016), *vasa-GFP-UTR-vasa* (291 – Wolke et al., 2002); *PA-GFP-UTR-globin* or *cd18-UTR-nanos3* were used as control RNA (A918 or 554, respectively – Hartwig et al., 2014). To generate *dn-rock-UTR-globin* construct, zebrafish *rock2* lacking the C-terminus (Marlow et al., 2002) was cloned upstream of *Xenopus globin* 3’UTR. For *ca-mlck-UTR-globin*, zebrafish *mlck* open reading frame lacking the autoinhibitory domain and the calmodulin binding site (Blaser et al., 2006) was cloned upstream of *Xenopus globin* 3’UTR. Morpholinos used (ZFIN names in parenthesis): *cxcr4b* MO (MO1-cxcr4b), *cxcl12a* MO (MO4-cxcl12a), *sox32* MO (MO2-sox32). For details on the RNA and morpholinos used in each experiment, see Table S1.

### Dextran gel preparation

To introduce artificial physical barriers into live embryos, we injected methacrylated dextran (DexMA) hydrogels. These hydrogels are protein-adsorption-resistant, which renders them biologically inert (Trappmann et al., 2017). Due to their non-swelling nature, the hydrogels retain their initial shape within tissues and are therefore especially suitable for generating barriers *in vivo*. In brief, a solution containing DexMA (71% methacrylation, 4.4% (w/v)), varying amounts of a crosslinker peptide (sequence CGPQGIAGQGC; GenScript, Piscataway, NJ, USA) and FluoSpheres 100 nm Yellow Green fluorescent beads (0.01% of gel volume; Invitrogen, Waltham, MA, USA) in PBS was prepared. The pH was adjusted to 8.0 using 1M NaOH to initiate hydrogel gelation, and the pre-gel solution was immediately injected into the interstitial space of zebrafish embryos. Final crosslinker concentration of 25.2 mM yielded soft hydrogels (Young’s modulus of 0.5 +/− 0.1 kPa), whereas stiff hydrogels (Young’s modulus of 4.1 +/− 0.4 kPa) were produced using 40.4 mM concentration. Importantly, changing the concentration of crosslinker does not cause change in the porosity of the gel (Trappmann et al., 2017).

### Analysis of PGC response to dextran gels

Embryos were injected at the early 1-cell stage, then incubated in 28°C. When the embryos reached 3 hpf, the DexMA gel was injected into the interstitial space. The embryos were then incubated at 25°C. At 7 hpf, the embryos that retained the gel were selected for, dechorionated and imaged using a spinning disc confocal microscope (Visitron Systems, Puchheim, Germany) with a 20x magnification water-immersion objective, capturing 100 μm Z-stack with 10 μm step size for 1 h at 30-s time intervals. The movies were then analyzed using the Fiji software (Schindelin et al., 2012). To correct for gastrulation tissue movement, the movies were registered based on the imaging channel used for detecting the gel employing the Correct 3D drift function. For the PGCs that came into contact with the gel during the time lapse acquisition, an interval of 6 min was set starting 1 min before the encounter. In addition, a control cell that did not come into contact with the gel was chosen. Then, based on the change in the front actin polarization during the 6-minute interval (1-minute prior and 5-minute after interaction with the gel), the response of both cells was scored.

To calculate response time, cells were followed after encountering the gel until turning or polarity loss responses occured, regardless of the 6-minute interval.

### Generation of single-germ-layer embryos

For the conversion of embryonic cells into mesodermal tissue, wild-type AB or transgenic females were crossed with *Tg(−1.8gsc:GFP)* males. The embryos were injected at the 1-cell stage and kept at 28°C until imaging. Unless mentioned otherwise, half an hour prior to imaging, the embryos were selected for strong and uniform expression of the GFP signal, dechorionated and prepared for microscopy.

For the conversion of embryonic cells into ectodermal tissue, *MZoep^tz257^* fish that contained the transgenes as indicated in Table S1 were incrossed. The embryos were injected at the 1-cell stage, and were then kept at 28°C. Prior to imaging, the embryos were dechorionated and prepared for microscopy.

### Expression of somatic RNAs in PGCs and in cells of the different germ layers

Embryos were fixed at 10 hpf in 4% PFA for 2 h at room temperature (RT). Then, the RNAscope procedure was performed as in (Gross-Thebing et al., 2014). The same probes as in (Gross-Thebing et al., 2020) were used (Biotechne, Minneapolis, MN, USA): *sox2* (catalog no. 494861-C3) and *tp63* (catalog no. 475511-C3) to label ectoderm derivatives; *sox17* (catalog no. 494711-C3) to label endodermal cells; and *pcdh8* (catalog no. 494741-C3), *pax8* (catalog no. 494721-C3), and *ntla* (catalog no. 4835511-C2) to label mesodermal cells. The PGCs were imaged on a confocal microscope (Zeiss, Oberkochen, Germany), using 40x magnification water-immersion objective, acquiring 24 μm Z-stacks with 2 μm step size).

### Locating PGCs in converted embryos

Embryos were converted into ectoderm or mesoderm and coinjected with *gfp-FTASE-UTR-globin*, fixed at 8 hpf in 4% PFA for 2 h at RT and washed in PBS. Embryos were imaged on a confocal microscope (40x water-immersion objective, 100 μm Z-stack with 5μm step size).

### Expression of germline RNA markers in PGCs

Converted and control embryos were fixed at 8 hpf in 4% PFA overnight at 4°C. The RNAscope procedure was then performed as in (Gross-Thebing et al., 2014). The following probes (Bio-Techne, Minneapolis, MN) were used: *nanos3* (catalog no. 404521-C2) and *vasa* (catalog. No. 407271-C3). Embryos were imaged on a confocal microscope using a 63x magnification water-immersion objective, acquiring Z-stacks with a 2 μm step size that include the entire PGC (as determined by the *nanos3* probe). The Z-stacks were projected with sum slices method (Fiji), and the average intensity of the *nanos* probe within the cell area was measured (the cell area was determined by the distribution of the *nanos3* probe). Background signal intensity was measured at two locations adjacent to the PGCs, and the average background intensity was subtracted from the average cell intensity. The same procedure was performed for the *vasa* probe channel.

### Tracking of PGCs located within the converted tissues

Embryos were converted into mesoderm or ectoderm as described above. Before imaging, the embryos were ramped in 1% low-melting agarose, and imaged between 7 and 8 hpf using a spinning disc confocal microscope (10x water-immersion objective, 250 μm Z-stack with 10 μm step size, 1 h with 2-min time interval). PGCs and somatic-cell nuclei (labeled with H2A-tagBFP) were tracked using the Imaris software (Oxford Instruments, Abingdon, UK), where the movement of PGCs relative to that of somatic nuclei was followed.

### Analysis of blebbing activity

PGCs were imaged using a spinning disc confocal microscope between 8 and 10 hpf (40x water-immersion objective, 11-min time lapses with 5-s time interval) and cells that were moving forward for longer than 3 min were included in the analysis. To quantify blebbing frequency, blebbing events were counted and divided by the analyzed time period (in minutes). The area of each bleb was measured using the freehand selection tool in Fiji at the time of maximum expansion and this value was divided by the cell area measured at the same time point to provide the “bleb size” parameter. Blebbing frequency and average bleb size values were normalized to the average value measured in mesoderm conversion.

### Analysis of nuclei density

Converted embryos were fixed at 8 hpf in 4% PFA for 2 h at RT, washed with PBS and stained with Hoechst (1:10000 in PBS + 0.1% Tween, Thermo Fisher Scientific, Waltham, MA, USA) overnight at 4°C. After washing in PBS, the embryos were ramped in 1% low-melting agarose drop and imaged using confocal microscope (40x objective, Z-stacks with 2μm step size that include the signal from external EVL cells to the yolk). The nuclei were segmented in 3D using the Imaris software and average distance to 5 closest neighbors was computed as a measure of cell density.

### E-cadherin staining

Converted embryos were fixed at 8 hpf in 4% PFA for 2 h at RT, washed with PBS + 0.1% Tween and transferred to Dent’s solution (20% DMSO, 80% methanol). E-cadherin immunostaining was performed as previously described (Blaser et al., 2006). To detect E-cadherin in zebrafish, mouse polyclonal anti-E-cadherin (cat. no. 610181, BD Bioscences, Franklin Lakes, NJ; internal no. P22) was used at 1:100 dilution. For labeling, goat anti-mouse secondary antibody conjugated with Alexa Fluor 568 (cat. no. A-11031, Thermo Fisher Scientific; internal no. S33) was used at 1:1000 dilution. Nuclei were stained with Hoechst (1:10000). Embryos were imaged on the confocal microscope, acquiring Z-stacks with 2 μm step size. Starting from the layer of EVL cells, 10 consecutive slices were projected using the *Average intensity projection* function in Fiji. The mean intensity of the fixed size area was measured at the centre of Z-projection. To control for the unspecific binding of the secondary antibody, control embryos were subject to the immunostaining procedure without adding the primary antibody. The average intensity value measured in such control embryos was substracted from the values measured for stained embryos. Then, for each repeat, the values were normalized to the average value of mesoderm embryos obtained in this repeat.

### PGC transplantation

Donor and host embryos were injected (host embryos were 2 h younger). When the donor embryos were about to reach 5 hpf and host embryos 3 hpf, the embryos were dechorionated. PGCs from donor embryos were located under the UV-fluorescent binocular (Leica, Wetzlar, Germany) and transplanted using a 50-μm wide transplantation needle (BioMedical Instruments, Zoellnitz, Germany) into the host embryos. Following the transplantation, the host embryos were incubated at 31°C until they reached the 8 hpf stage. Between the 8 and 10 hpf stages, the PGCs that were surrounded by the host cells (marked with mCherry-H2B) were imaged using a spinning disc confocal microscope (40x water-immersion objective, 10 μm Z-stack with 5-μm step size, 11 min with 5-s time interval). The images were subjected to maximum intensity projection, and blebbing behavior was analyzed as described above. Individual blebbing frequency and average bleb size values were then normalized to the average value in wild-type host cells (Fig. S2) or control host cells (Fig. 2D, G). In case of transplantations into mesoderm converted embryos (Fig. 2G), host embryos were 0.5 h older than the donor embryos. Before imaging, host embryos were selected for uniform *gsc:GFP* expression.

For tracking (Fig. S4), transplanted embryos were imaged between 8 to 9 hpf using a spinning disc confocal microscope (10x water-immersion objective, 200 μm Z-stack with 10 μm step size, 1 h with 2-min time interval). PGCs and somatic cells were then tracked as described above. For each repeat, speed and track straightness of cells were normalized to the average speed and track straightmess value of control cells obtained on that experimental day.

### FRET measurements

Mesoderm-converted embryos were selected based on their morphology (i.e., cells remaining at the animal pole of the embryos). Between 8 and 10 hpf, the germ cells were imaged using a confocal microscope (40x water-immersion objective, for at least 3 min at 8-s interval) and analyzed as previously described (Tarbashevich et al, 2015). For each repeat, whole cell FRET values were normalized to the average FRET value of PGCs located within mesodermal tissue obtained on the same day. In addition, in the RhoA FRET experiment, each cell was divided into four sections along the longest axis, with the front and back defined as first and last quarter of the cell. The front/back FRET ratios were then calculated.

### Ezrin distribution analysis

Mesoderm-converted embryos were selected based on their morphology. Between 8 and 10 hpf, the germ cells were imaged using a spinning disc confocal microscope (40x water-immersion objective, 10 μm Z-stack with 5-μm step size, 11 min with 5-s interval). For each cell, 20 consecutive frames of a polarized, forward-moving cell were selected. Using the Fiji software, the cell cortex was segmented in the following manner: Based on lifeact signal, the cell was thresholded and converted into a binary mask. After applying the *Fill holes* function, a copy of the cell mask was subjected to the *Erode* operation four times. Using the *Image-calculator* function, the difference between the original cell mask and the eroded cell mask was generated and used as a cortex mask. Ezrin signal was masked with the cortex mask and analyzed with a custom Python script. The center of the cell was determined as the center of mass of a binarized image of the selected contour. Then, for each image in the stack, the contour was split in 120 sections of *θ* = 3° wide, and all intensity values within a section were averaged. A Gaussian function of the form:

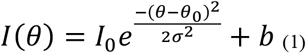

was then fitted to the obtained intensity distribution, where σ is the width of the peak in the fluorescence distribution. Finally, for plotting, the data were normalized by the amplitude and shifted so that the peak was centered on 180°.

### Actin-rich protrusion time measurements

Embryos were converted into specific germ layers and incubated at 28°C. Between 8 and 10 hpf, the PGCs were imaged using a spinning disc confocal microscope (63x water-immersion objective, 4 min with 500-ms interval). Movies were aquired at a single focal plane that included the front and the back of the cell. We counted the frames that contained the actin-rich front of the forward-moving cell and did not show blebs. Then, we divided that value by the number of frames in which the cell was polarized and moved forward.

### Tracking of PGCs exhibiting increased blebbing

Converted embryos were ramped at 8 hpf in 1% low-melting agarose and imaged on spinning disk confocal microscope (10x water-immersion objective, 200 μm Z-stack with 10 μm step size, 1 h with 2-min time interval). PGCs and somatic cells were tracked as described above. For each repeat, speed and track straightness of cells were normalized to the average speed and track straightness value of respective control cells obtained on that experimental day.

### Methodology, statistics

All the experiments were repeated at least three times, except for the E-cadherin staining (Fig. S6), which was done twice. For each experiment, embryos from the same egg clutch were allocated to a particular treatment group blindly. Investigators were not blinded during the data acquisition or during the analysis, except for the analysis of the hydrogel experiments (Fig. 4E-G and Fig S7), which was double blinded. Before choosing statistical test, data was tested for normal distribution using D’Agostino-Pearson test. Normally distributed data was subjected to t-test, while non-normal distribution was subjected to Mann-Whitney test. In cases where normally distributed data had unequal variances, t-test with Welsch’s correction was employed.

## Acknowledgements

We would like to thank E.-M. Messerschmidt, I. Sandbote and U. Jordan for technical help, J. Wegner for help in preparing schematics and C. Brennecka for critical reading of the manuscript.

## Author contributions

Conceptualization: L.T., E.R; Methodology: L.T., D.B., H.L., B.T.; Software: B.E.V.; Formal analysis: L.T.; Investigation: L.T., D.B., K.T.; Resources: B.E.V.; B.T.; Data curation: L.T., B.E.V.; Writing – original draft: L.T., E.R.; Writing – review and editing: H.L., B.E.V.; B.T.; Visualization: L.T.; Supervision: B.T., E.R.; Funding acquisition: B.T., E.R.

