## Supplemental Information for "Primordial germ cells adjust their protrusion type while migrating in different tissue contexts *in vivo*"

### Supplementary information
for:

Primordial germ cells adjust their protrusion type while migrating in different tissue contexts *in vivo*

Lukasz Truszkowski, Dilek Batur, Hongyan Long, Katsiaryna Tarbashevich, Bart E. Vos, Britta Trappmann, Erez Raz

Contents:

Supplementary Figure S1-S7
Movie 1-6 titles and captions

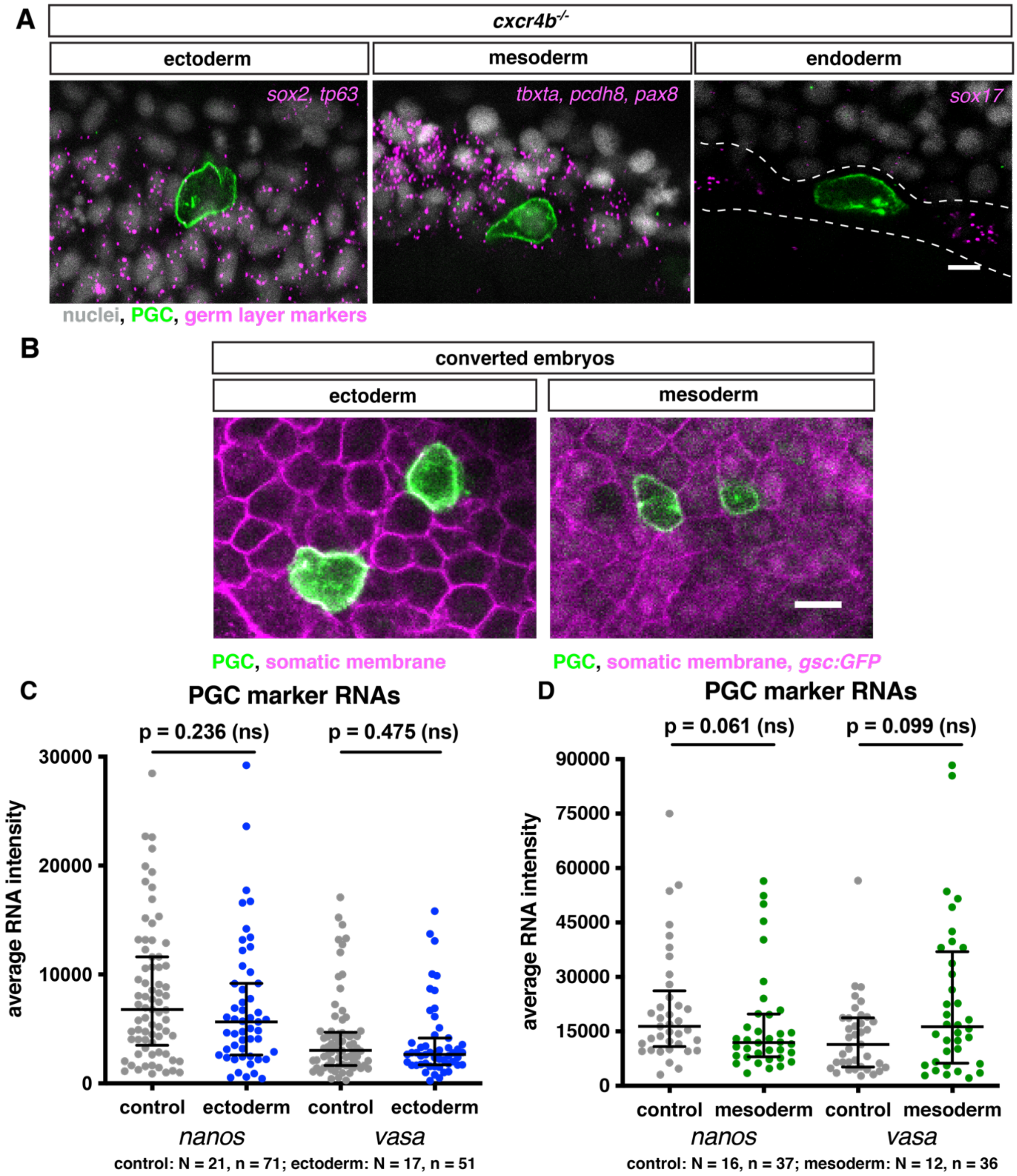


**Figure S1. Preservation of PGC identity in different germ layers.** (A), Representative PGCs lacking guidance cues (green) located within tissues of ectodermal, mesodermal and endodermal origin of wild-type embryos (magenta). The thin endodermal layer is marked with a white dashed line. Scale bar, 20 µm. (B) Examples of PGCs (green) residing among the somatic cells of embryos whose cells were converted into ectoderm or mesoderm (left and right panels respectively, magenta). Scale bar, 20 µm. (C-D), Quantification of *nanos* and *vasa* RNA expression levels in PGCs located within control or ectoderm-converted embryos (C) and control or mesoderm-converted embryos (D). For both graphs, lines represent median value +/- interquartile range (IQR). Mann-Whitney test was performed. N and n represent numbers of embryos and cells, respectively.

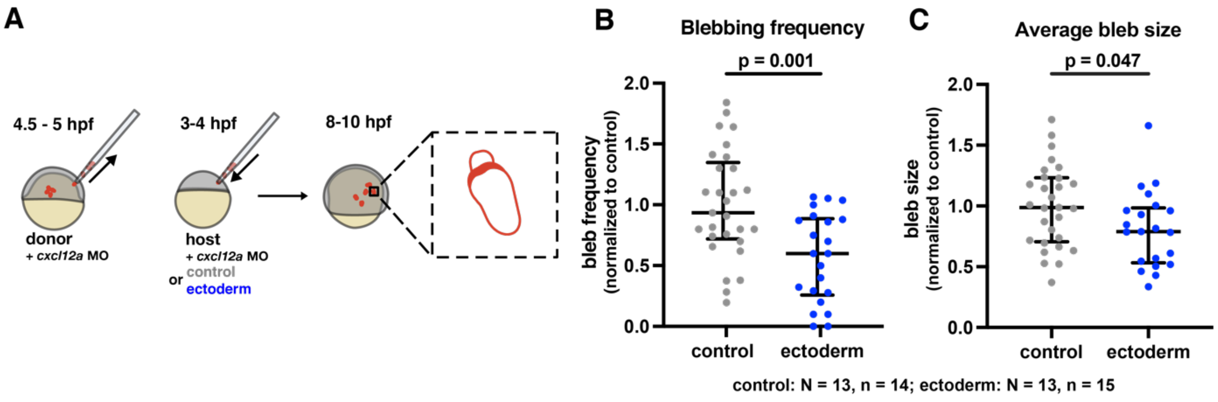


**Figure S2. PGC response to different environments.** (A), Illustration of the PGC transplantation experiment. Labeled wild-type PGCs were transplanted into control and ectoderm-converted embryos. After 4 to 6 h, the blebs produced by the transplanted cells were analyzed. Control embryos were wild-type embryos. (B-C), Bleb formation frequency (B) and average bleb size (C) in transplanted PGCs. Lines show median +/- IQR. Mann-Whitney test was performed. N and n represent numbers of embryos and cells, respectively.

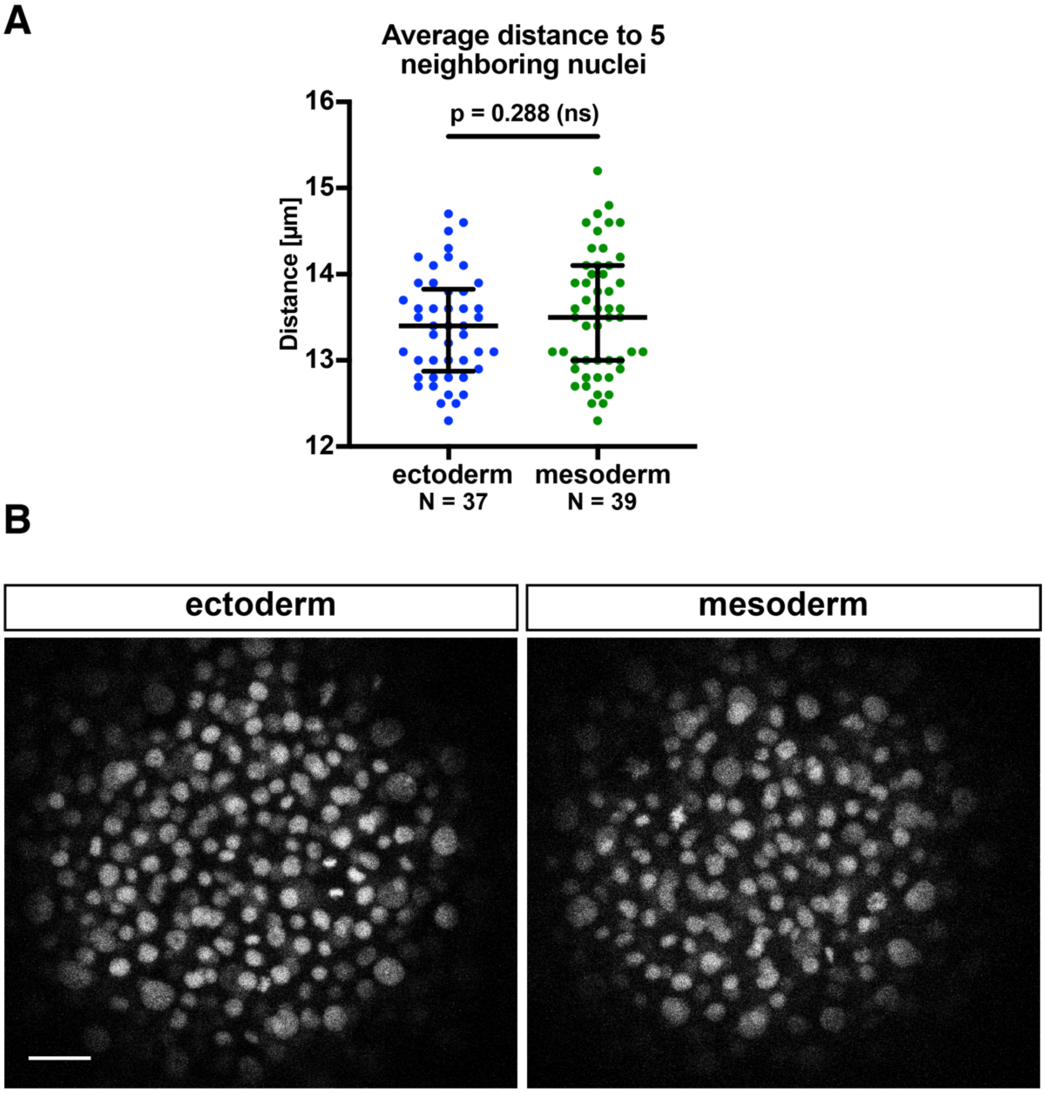


**Figure S3. Cell density in converted embryos.** (A) Average distance of somatic cell nuclei to 5 neighboring nuclei in converted ectoderm and mesoderm tissues. Lines show median +/- IQR. Ectoderm: Two tailed t-test was performed. N represents number of embryos. (B) Representative images of converted embryos with Hoechst-stained nuclei. Scale bar, 30 µm.

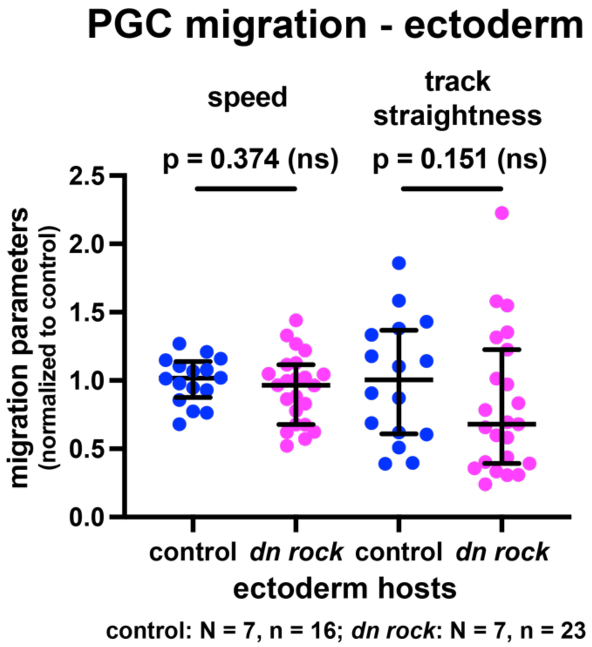


**Figure S4. PGC migration features in modified ectodermal environment.** Migration speed and track straightness of PGCs transplanted into ectoderm-converted embryos injected at the 1-cell stage with control RNA or *dn rock* RNA (performed as presented in Fig. 2D). Lines show median +/- IQR. Two-tailed t-test was performed for speed comparison, Mann-Whitney test – for track straightness comparison. N and n represent numbers of embryos and cells, respectively.

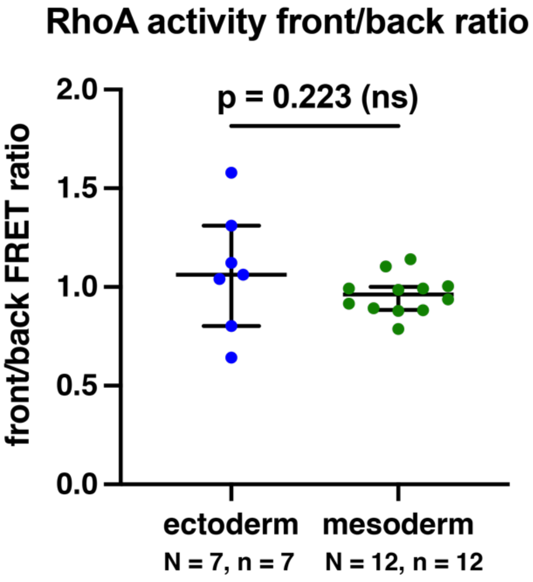


**Figure S5. Distribution of RhoA activity in PGCs migrating within different cellular environments.** Ratio between the RhoA FRET levels at the front and the back of the cell. Lines show median +/- IQR. Two-tailed t-test was performed. N and n represent numbers of embryos and cells, respectively.

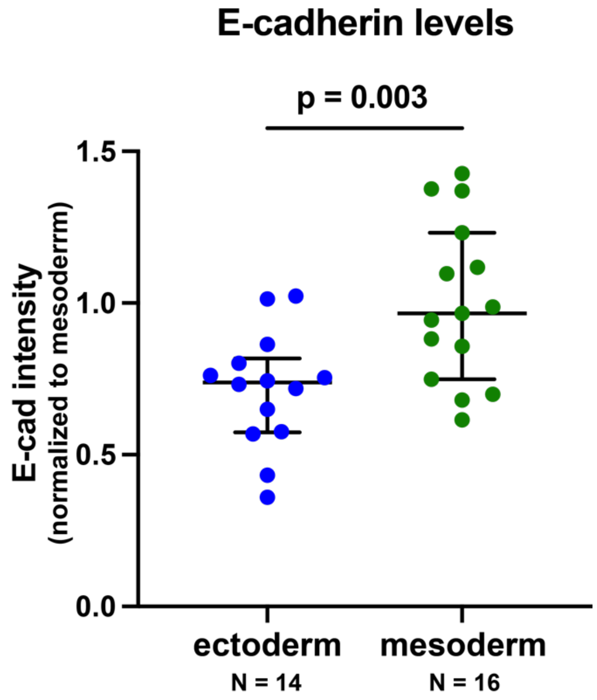


**Figure S6. E-cadherin levels in converted embryos.** E-cadherin antibody staining in ectoderm- and mesoderm-converted embryos. Lines show median +/- IQR. Two-tailed t-test was performed. N represents number of embryos.

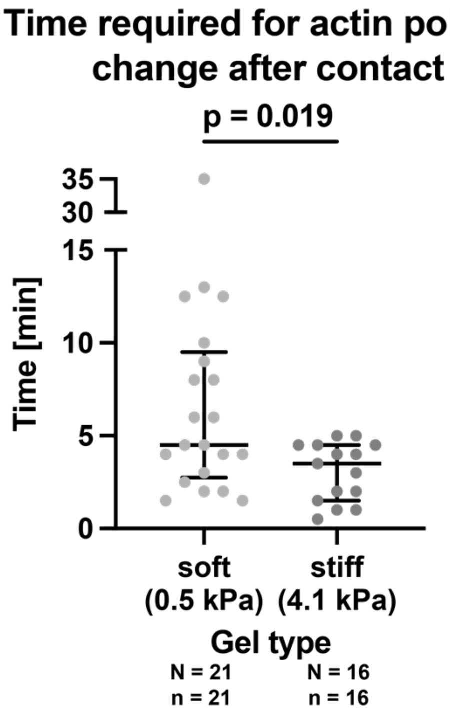


**Figure S7. Time required for a reaction.** Plot of the time required for PGCs to change polarity upon contact with the gel, regardless of the interval used in Fig. 4G. Mann-Whitney test was performed. N and n represent numbers of embryos and cells, respectively.

**Movie titles and captions**

**Movie 1. A migrating PGC that generates actin-rich protrusions.** Cell was labelled with lifeact-mCherry. Snapshots from the movie are presented in upper panels of Fig 2J.

**Movie 2. A migrating PGC that generates blebs.** Cell was labeled with lifeact-mCherry. Snapshots from the movie are presented in lower panels of Fig 2J.

**Movie 3. Protrusions in a polarized and an unpolarized cell located within ectodermal environment.** 3D projection of Z-stack the includes the whole cell (28 µm stack, with 2µm step size). Cells were labeled with lifeact-mCherry. Top view, a migrating, polarized cell (left), and an apolar cell generating blebs in different directions (right). Side view, with actin enrichments segmented using the "Surfaces" function of Imaris. Actin-rich pseudopods and blebs that are clearly visible in both views are marked with plus (+) and asterisk (*) symbols, respectively. The segmented actin enrichment is always present at the front of pseudopods, in contrast to blebs, in which actin is absent during the initiation and inflation phases. Both types of protrusions are three-dimensional.

**Movie 4-6. PGC response to contact with the gel.** Time 0 shows the estimated moment that the cell (magenta) contacts the gel (green). Snapshots from movies are used in Fig.4F to present a turning response (Movie 4), loss of polarity (Movie 5) and no change in polarity (Movie 6).
